# Neutrophils use selective autophagy receptor p62/SQSTM1 to target *Staphylococcus aureus* for degradation *in vivo* in zebrafish

**DOI:** 10.1101/604884

**Authors:** Josie F Gibson, Tomasz K Prajsnar, Christopher J Hill, Amy K Tooke, Justyna J Serba, Rebecca D Tonge, Simon J Foster, Andrew J Grierson, Philip W Ingham, Stephen A Renshaw, Simon A Johnston

## Abstract

Autophagy leads to degradation of cellular components and has an important role in restricting intracellular pathogens. Autophagy receptors, including p62, target invading intracellular pathogens to the autophagy pathway for degradation. *Staphylococcus aureus* is a significant pathogen of humans and often life-threatening in the immunocompromised. Increasing evidence demonstrates that *S. aureus* is an intracellular pathogen of immune cells and may use neutrophils as proliferative niche but the intracellular fate of *S. aureus* following phagocytosis by neutrophils has not previously been analysed *in vivo. In vitro*, p62 is able to co-localise with intracellular *Staphylococcus aureus*, but whether p62 is beneficial or detrimental in host defence against *S. aureus in vivo* had not been determined.

Here we use zebrafish to determine the fate and location of *S. aureus* within neutrophils throughout infection. We show that Lc3 and p62 recruitment to phagocytosed *S. aureus* is altered depending on the bacterial location within the neutrophil. We also show rapid Lc3 marking of bacterial phagosomes within neutrophils which may be associated with subsequent bacterial degradation. Finally, we find that p62 is important for controlling cytosolic bacteria demonstrating for the first time a key role of p62 in autophagic control of *S. aureus* in neutrophils.

## Introduction

Autophagy (macroautophagy) is a process of cellular self-degradation, in which damaged or redundant cellular components are taken into an autophagosome and subsequently trafficked to the lysosome for degradation; these degraded components can then be recycled for alternative uses by the cell (Mizushima *et al.*, 2008; Tanida, 2011). During infection, autophagy is used by host cells to degrade invading pathogens, a process termed xenophagy (Gatica, Lahiri and Klionsky, 2018; Sharma *et al.*, 2018).

Autophagy is considered largely non-selective of the cargo to be degraded, classically being induced by starvation conditions. However, selective autophagy is a process that enables specific cargo to be directed into the autophagy pathway, which can be used to target invading pathogens. Selective autophagy uses autophagy receptors (ARs), proteins that interact with both autophagy machinery and the cargo to be degraded (Popovic and Dikic, 2012; Rogov *et al.*, 2014). Many ARs are involved in targeting invading pathogens, including p62 (also named sequestosome 1 (SQSTM1)), neighbour of Brca1 gene (NBR1), optineurin (OPTN) and nuclear dot protein 52 (NDP52) (Farré and Subramani, 2016).

Loss of autophagy function, for example through mutations in key autophagy genes, can increase the risk of infection with intracellular pathogens (Levine, Mizushima and Virgin, 2011). It is well established that pathogen presence can induce host cell autophagy and that pathogens can be degraded by this pathway. Intracellular pathogens such as *Mycobacterium marinum, Shigella flexneri* and *Listeria monocytogenes* (Mostowy *et al.*, 2011; Zhang *et al.*, 2019) can be targeted by ARs for degradation. Conversely, pathogens have evolved to be able to block or subvert immune defences, and autophagy is no exception. Indeed, many bacterial pathogens are able to inhibit induction of autophagy or to reside within the autophagy pathway by preventing lysosomal fusion, or even avoid making any contact with autophagic machinery (Deretic and Levine, 2009). In some cases it is beneficial to the pathogen to up-regulate the autophagy pathway, for example *Legionella pneumophila, Coxiella burnetii* and *Salmonella enterica* serovar typhimurium (Hernandez *et al.*, 2003; Amer and Swanson, 2005; Gutierrez *et al.*, 2005). The outcome of host-cell autophagy therefore differs between various invading pathogens.

*Staphylococcus aureus* is a bacterial pathogen that is able to reside within neutrophils as an intracellular niche (Thwaites and Gant, 2011; Prajsnar *et al.*, 2012). Autophagy has been implicated in *S. aureus* infection but there are conflicting reports suggesting autophagy might be either beneficial (Schnaith *et al.*, 2007) or detrimental for *S. aureus* (Neumann *et al.*, 2016). Intracellular pathogens, including *S. aureus*, are able to escape the phagosome into the cytosol (Bayles *et al.*, 1998), likely through toxins secreted by the bacteria or membrane rupture due to bacterial growth. Once in the cytosol, bacteria can be ubiquitinated and targeted by ARs (Farré and Subramani, 2016). Indeed, p62 in fibroblasts and epithelial cells has been shown to localise to cytosolic *S. aureus* leading to autophagsome formation *in vitro* (Neumann *et al.*, 2016; Singh *et al.*, 2017). Therefore, we investigated whether p62 recruitment is employed by neutrophils in *S. aureus* infection and what influence selective autophagy has on infection outcome *in vivo*.

In order to examine the role of neutrophil autophagy in *S. aureus* infection, we compared the fate of bacterial cells following Lc3 and p62 recruitment. We tested the role of p62 in pathogen handling *in vivo*, using the genetic tractability of the zebrafish to create a neutrophil-specific p62-GFP transgenic reporter, and a *p62* activity deficient mutant. With this approach we show that p62 is recruited to cytosolic *S. aureus* and disruption of p62 expression or function adversely affects *S. aureus* infection outcome.

## Results

### *Staphylococcus aureus* location within neutrophils changes throughout infection

Autophagy responses have been demonstrated to change throughout the progression of infection. Targeting of pathogens by autophagy receptors is likely to occur at later time points in infection. Therefore, to determine the fate and location of *S. aureus* in neutrophils during infection, *S. aureus* expressing mCherry was inoculated and imaged at early (2 to 5 hours post infection (hpi)) and late (24 to 28hpi) time points. Initially, the well-established *Tg(mpx:eGFP)i114* line that specifically marks neutrophils with EGFP (Renshaw *et al.*, 2006) was used to analyse the fate of intracellular *S. aureus* throughout infection. Imaging throughout the whole organisms demonstrated a marked reduction in the number of bacterial cells within individual neutrophils), and that the number of neutrophils containing *S. aureus*, between 2 and 24 hours post infection (Figure 1A, B). This suggested to us that neutrophils were able to degrade intracellular *S. aureus* effectively throughout infection. Indeed, video timelapse of *Tg(mpx:eGFP)i114* larvae infected with mCherry *S. aureus* demonstrated that bacteria can be effectively degraded by the host neutrophils (Figure 1C), although in other cases the bacterial infection is not controlled (Supplementary Figure 1A).

**Figure 1:**
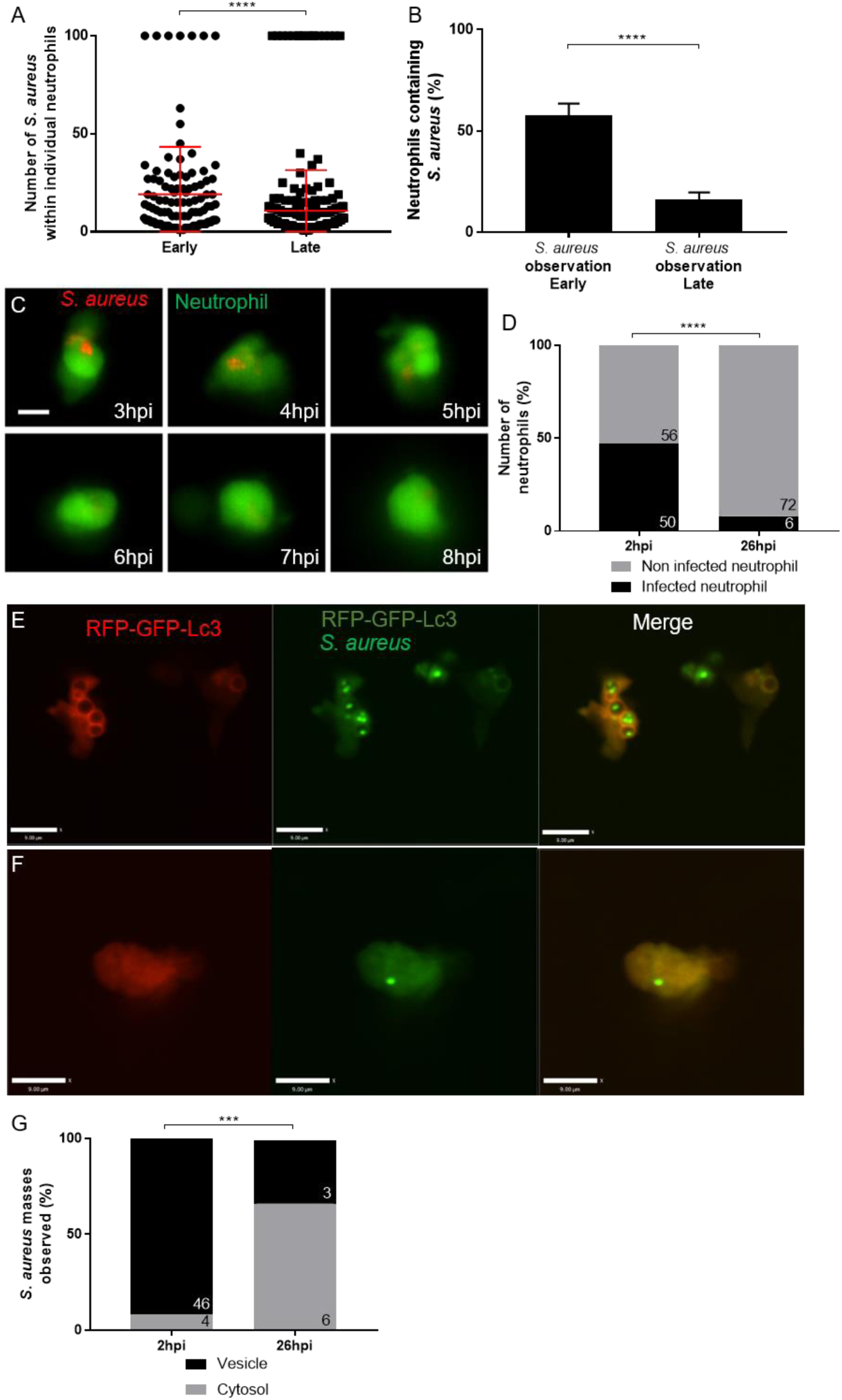
*Staphylococcus aureus* location within neutrophils changes from vesicular to cytosolic throughout infection. **A-C** *Tg(mpx:eGFP)*i114 larvae were injected at 1dpf with 1500cfu SH1000-mCherry *S. aureus*, and imaged at early (1-5hpi) and late (24-28hpi) time points **A** Number of bacteria contained in neutrophils, with maximum 100 bacterial cells counted, (whole larvae imaged, n=11-13, Mann-Whitney test, ****p<0.0001, +/- SD) **B** Proportion of neutrophils containing bacteria (whole larvae imaged, n=11-12, unpaired t-test, ****p<0.0001, +/- SEM) **C** *Tg(mpx:eGFP)*i114 larvae were injected at 1dpf with 1500cfu SH1000-mCherry *S. aureus*, and imaged at 3 hours post infection. Images were captured every 5 mins for 12 hours at multiple z planes to follow infected neutrophils over time. Scale bar is 5 μm **D-G** *Tg(lyz:RFP-GFP-Lc3)*sh383 larvae were injected at 2dpf with GFP *S. aureus*, and imaged in the CHT at 2hpi, and ∼26hpi. **D** The proportion of infected or non-infected neutrophils at 2hpi and 26hpi (****p<0.0001 Chi Square test, n=3, 17 2hpi larvae, 11 26hpi larvae) **E** *S. aureus* with Lc3 marking the entire vesicle, scale 9um, demonstrating a vesicle **F** *S. aureus* in the cytosol, scale 9 μm **G** Proportion *S. aureus* events observed within vesicles or cytosol at 2hpi and 26hpi (***p<0.001, Fisher’s exact test, n=3, 17 larvae at 2hpi, and 11 larvae at 26hpi).

We next sought to determine the location of bacteria, and relationship to autophagic machinery, within neutrophils. To do this we used the newly generated *Tg(lyz:RFP-GFP-Lc3)sh383* (Prajsnar *et al.*, 2019). We first confirmed that, in the caudal hematopoietic tissue (CHT), the infection dynamics were similar to the *Tg(mpx:eGFP)i114* line, with a significant reduction in intracellular bacteria by 26hpi, indicating bacteria are efficiently controlled and a significant reduction in infected neutrophils was observed (Figure 1D). Importantly, the number of neutrophils analysed in the CHT, used for analysis throughout this study, did not significantly change between 2dpf and 3dpf (Supplementary Figure 1B), demonstrating that the change in proportions of infected neutrophils is not due to a large increase in neutrophil number between these time points. The labelling of *S. aureus* containing vesicles enabled the identification of intracellular bacteria that were within a vesicle (Figure 1E) or free in the cytosol (Figure 1F), as well as non-labelled vesicles, or vesicles marked with Lc3 puncta (Supplementary Figure 1C, D). We found that the proportion of bacteria within vesicles was significantly reduced over time post-injection, whereas the number of bacteria within the cytosol remains relatively constant at a low level, despite becoming proportionally higher relative to vesicular bacteria (Figure 1G). Thus, *S. aureus* phagocytosed by a neutrophil are initially located in a phagocytic vesicle and are subsequently degraded. However, a smaller proportion of *S. aureus* is able to survive to later infection time points, and these predominantly reside in the cytosol.

### Generation and characterisation of an *in vivo* neutrophil GFP-p62 reporter line

A previous study identified colocalisation of p62 with *S. aureus* in non-immune cells (Neumann *et al.*, 2016). Our findings demonstrated a small but significant population of bacteria that were cytosolic, and therefore a possible target for p62 binding. Accordingly, we generated a transgenic neutrophil-specific p62 reporter zebrafish line to examine whether p62 and intracellular pathogens are co-localised *in vivo*. We used GFP fused via a small linker region to the N-terminus of *p62* in order to produce a fluorescently marked fusion protein expressed within neutrophils via the lysozyme C (*lyz*) promoter (Yang *et al.*, 2012). Using larvae with double labelled neutrophils, we were able to identify GFP expressing cells from the *Tg(lyz:eGFP-p62)i330* reporter line (hereafter called GFP-p62 reporter) also expressing mCherry (*Tg(lyz:nfsB-mCherry)sh260*) (Buchan *et al.*, 2019) in 98% of neutrophils observed (Supplementary Figure 2A-C).

We next examined whether the GFP-p62 protein is able to function as expected. Interestingly, in the double labelled larvae, GFP puncta but not mCherry puncta were seen (Supplementary Figure 2D). Similar p62 puncta that required UBD to function have been observed *in vitro* for endogenous p62 (Bjørkøy *et al.*, 2005). To test whether the GFP-p62 puncta observed in the GFP-p62 reporter line respond as expected, GFP-p62 reporter larvae were treated with autophagy inhibitor Bay-K8644 (known to block autophagy in zebrafish; (Williams *et al.*, 2008). As expected, there was a significant increase in the number of neutrophils which contained GFP-p62 puncta following Bay-K8644 treatment in comparison to non-treated controls (Supplementary Figure 2E), as well as a significant increase in the number of GFP-p62 puncta within individual neutrophils as expected for endogenous p62 (Supplementary Figure 2F). This suggests that the GFP-p62 puncta are not being processed through autophagy accumulate within the cell, as reported for endogenous p62 (Bjørkøy *et al.*, 2009).

As we had done for neutrophils and Lc3 positive vesicles, we examined the location of *S. aureus* throughout infection with our GFP-p62 reporter for consistency with *Tg(mpx:eGFP)i114* and *Tg(lyz:RFP-GFP-Lc3)sh383* (Figure 1). We found that there was a comparable reduction in the number of bacteria observed within neutrophils at 26hpi in comparison to 2hpi (Supplementary Figure 3A) and a reduction in the number of infected neutrophils from 2hpi to 26hpi (Supplementary Figure 3B). This suggested that neutrophils were efficiently degrading these bacteria, in agreement with Figure 1C.

Cytosolic bacteria are a possible target for p62 and *S. aureus* has previously been visualised within the cytosol of a neutrophil from murine infection studies (Gresham *et al.*, 2000).To identify *S. aureus* in the cytosol in our *in vivo* experiments in zebrafish we looked for regions of the cytosol that co-localised with S. aureus but without a reduction of GFP signal indicating a vacuole excluding the surrounding cytosol (containing GFP). We first confirmed we were able to clearly observe phagosomes containing bacteria with low GFP fluorescence consistent with *S. aureus* containing vacuoles, where host cell cytoplasm containing GFP, was excluded (p62GFP^low^, Supplementary Figure 3C). To further evidence this analysis, we determined that vesicles containing *S. aureus* visualised by TEM were empty of cellular components, in comparison to the cytosol (Supplementary Figure 3D), suggesting GFP^low^ areas represent vesicles. Finally, we looked for function differences consistent with the presence of a phagosomal membrane in GFP^low^ regions by examining pH differences using the pH sensitive dye pHrodo. We found examples of low pH in vesicles correlating with low cytoplasmic fluorescence (Supplementary Figure 3E), again suggesting GFP^low^ areas represent vesicles. Having characterised features consistent with a *S. aureus* containing vacuoles we were able to assign a subset of bacteria as being in either a damaged phagosome or located in the cytosol (p62GFP^high^, Supplementary Figure 3F); for the purpose of this study we are defining these bacteria as cytosolic as they are accessible to cytosolic proteins. We then assigned the cellular location of *S. aureus* by these features at 2hpi and 26hpi. We determined that the proportion of *S. aureus* within vesicles was significantly reduced by 26hpi (Supplementary Figure 3G), and that the number of bacteria within the cytosol is similar at both time points, in agreement with our *Tg(lyz:RFP-GFP-Lc3)sh383* data (Figure 1).

### Lc3 and p62 are recruited to *Staphylococcus aureus* within neutrophils

We determined that GFP-p62 puncta co-localise with *S. aureus* either marking a vesicle containing *S. aureus* (Figure 2A, supplementary video 1), or directly in contact with *S. aureus* located in the cytosol (Figure 2B, supplementary video 2). For puncta marking *S. aureus* in vesicles, no difference in the proportion of vesicles marked was observed at 2 or 26hpi, although the actual number of puncta marking was dramatically reduced by 26hpi (Figure 2C) as most bacteria had already been degraded. GFP-puncta marking bacteria in the cytosol were decreased at 26hpi (Figure 2D), as expected given that p62 is degraded with the cargo targeted for degradation (Bjørkøy *et al.*, 2009). We could show cytosolic GFP-p62 puncta were modulated by autophagy machinery-targeting drugs (Supplementary Figure 2E-F). In further agreement with this, comparison between infected and uninfected neutrophils showed there was no difference in the number of cytoplasmic GFP-p62 puncta at 2hpi but a significant reduction by 26hpi (Figure 2E, F), indicating these puncta are modulated by *S. aureus* infection. We next examined whether Lc3 is able to localise to vesicular and cytosolic *S. aureus*. At 2hpi and 26hpi there was no difference in the proportion of vesicles marked by Lc3, but most vesicular bacteria are degraded by 26hpi (Figure 2G), showing that a rapid Lc3 response to *S. aureus* infection occurs. In contrast, vesicles containing *S. aureus* are significantly more likely to have Lc3 puncta associated at 2hpi (Figure 2H, Supplementary Figure 1D), although most bacteria are still cleared by 26hpi and there was no significant change in the association of Lc3 puncta to *S. aureus* in the cytosol over time (Figure 2I).

**Figure 2:**
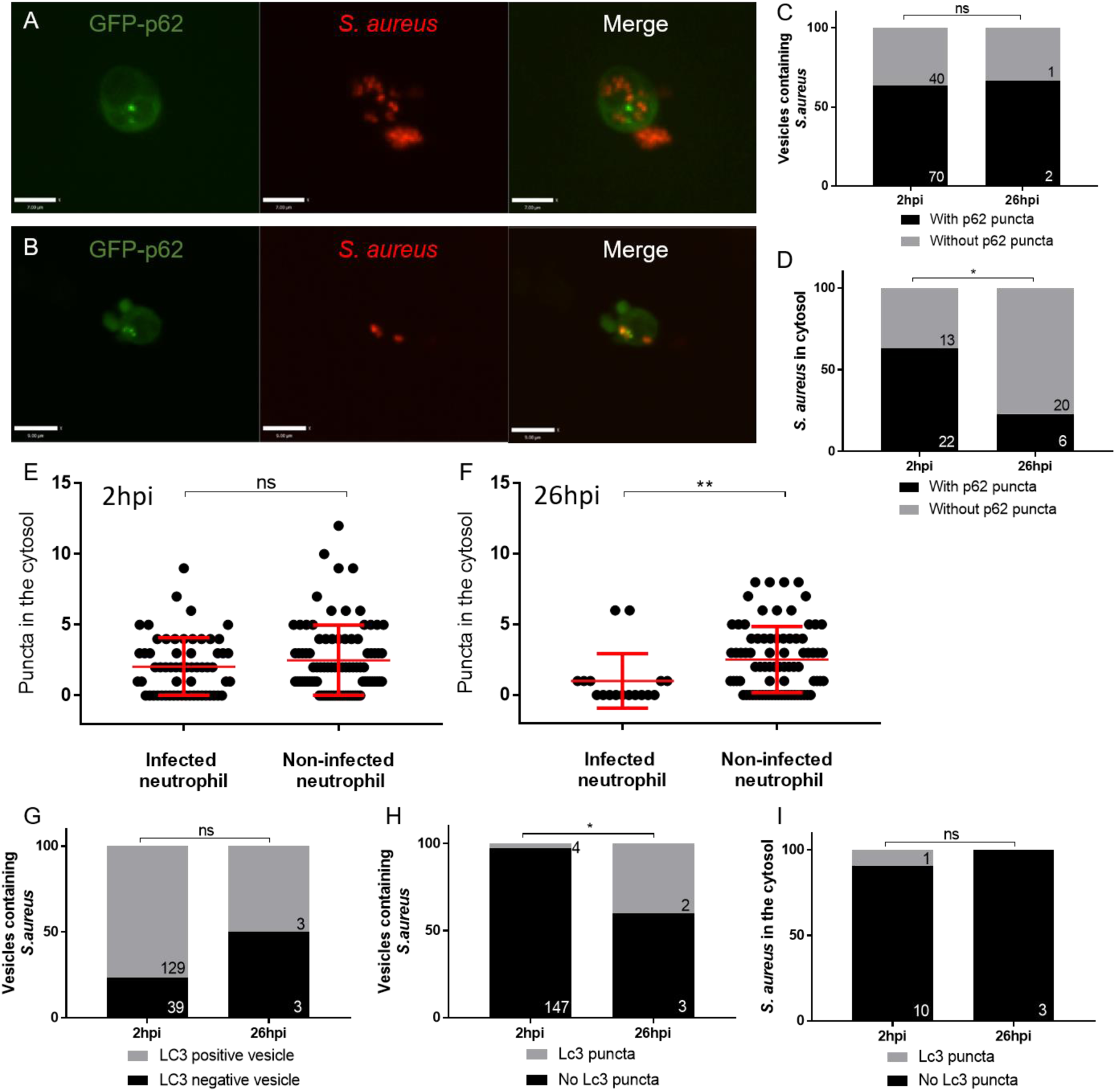
*In vivo* recruitment of GFP-p62 puncta during *S. aureus* infection. **A** Representative image of *S. aureus* observed within a likely “vesicle” with GFP-p62 puncta localisation, scale 7um **B** representative image of *S. aureus* observed within the cytosol with GFP-p62 puncta localisation, scale 9um **C** *S. aureus* within vesicles, co-localised with GFP-p62 at 2hpi and 26pi (CHT imaged, ns, Fisher’s exact test, n=3, 14 larvae at 2hpi, and 12 larvae at 26hpi) **D** *S. aureus* in the cytosol, co-localised with GFP-p62 at 2hpi and 26hpi (CHT imaged, *p<0.05, Fisher’s exact test, n=3, 14 larvae at 2hpi, and 12 larvae at 26hpi) **E** GFP-p62 puncta in the cytosol of infected and non-infected at 2hpi (CHT imaged, ns, Mann-Whitney test, n=3, error bars +/- SD, 14 larvae) **F** GFP-p62 puncta in the cytosol of infected and non-infected at 26hpi (CHT imaged, **p<0.01, Mann-Whitney test, n=3, error bars +/- SD, 12 larvae) **G-I** 2500cfu of GFP *S. aureus* injected into *Tg(lyzC:RFP-GFP-Lc3)sh383*, larvae imaged in the CHT at 2hpi (2hpi) and 26hpi (26hpi). **G** Lc3 association to the entire *S. aureus* vesicle at 2hpi and 26hpi (ns, Fisher’s test, n =3, 17 2hpi larvae, 11 26hpi larvae) **H** The number of *S. aureus* vesicles with Lc3 puncta (*p<0.05, Fisher’s test, n =3, 17 2hpi larvae, 11 26hpi larvae) **I** The number of *S. aureus* events in the cytosol with Lc3 puncta at 2hpi and 26hpi (ns, Fisher’s test, n =3, 17 2hpi larvae, 11 26hpi larvae).

### Loss of p62 reduces zebrafish survival following *Staphylococcus aureus* infection

We had demonstrated the steps of Lc3 and the autophagy receptor p62 recruitment *in vivo* in the degradation of *S. aureus* by neutrophils suggesting a function for p62 in immunity to *S. aureus* infection by targeting degradation of bacteria that escaped the phagosome. To test this prediction we examined the role of p62 in *S. aureus* zebrafish infection using a morpholino-modified antisense oligonucleotide (morpholino) targeting *p62* (van der Vaart *et al.*, 2014) to knock-down *p62* expression in the zebrafish larvae. Knock down of *p62* resulted in a significant reduction in zebrafish survival following *S. aureus* infection, compared to control larvae, supporting a requirement for *p62* in the control of *S. aureus* infection (Figure 3A). Knockdown of *p62* did not reduce larval survival for heat-killed *S. aureus* or the non-virulent but closely related bacterium *Micrococcus luteus* (Supplemental Figure 4B, C), suggesting p62 is important for restriction of pathogenic bacteria that escapes the phagosome. To further support this conclusion, we generated a *p62* mutant zebrafish (sh558) that lacked a functional UBD domain in *p62*, inhibiting the ability of p62 to bind to ubiquitinated cargo (Figure 3C). In agreement with our knock down study, the *p62* mutant zebrafish (sh558) larvae were significantly more susceptible to *S. aureus* infection than wild-type control zebrafish (Figure 3B). Thus, in addition to demonstrating how Lc3 and p62 were localised during intracellular handling of *S. aureus* by neutrophils, we could independently show the requirement of p62 in the outcome of infection.

**Figure 3:**
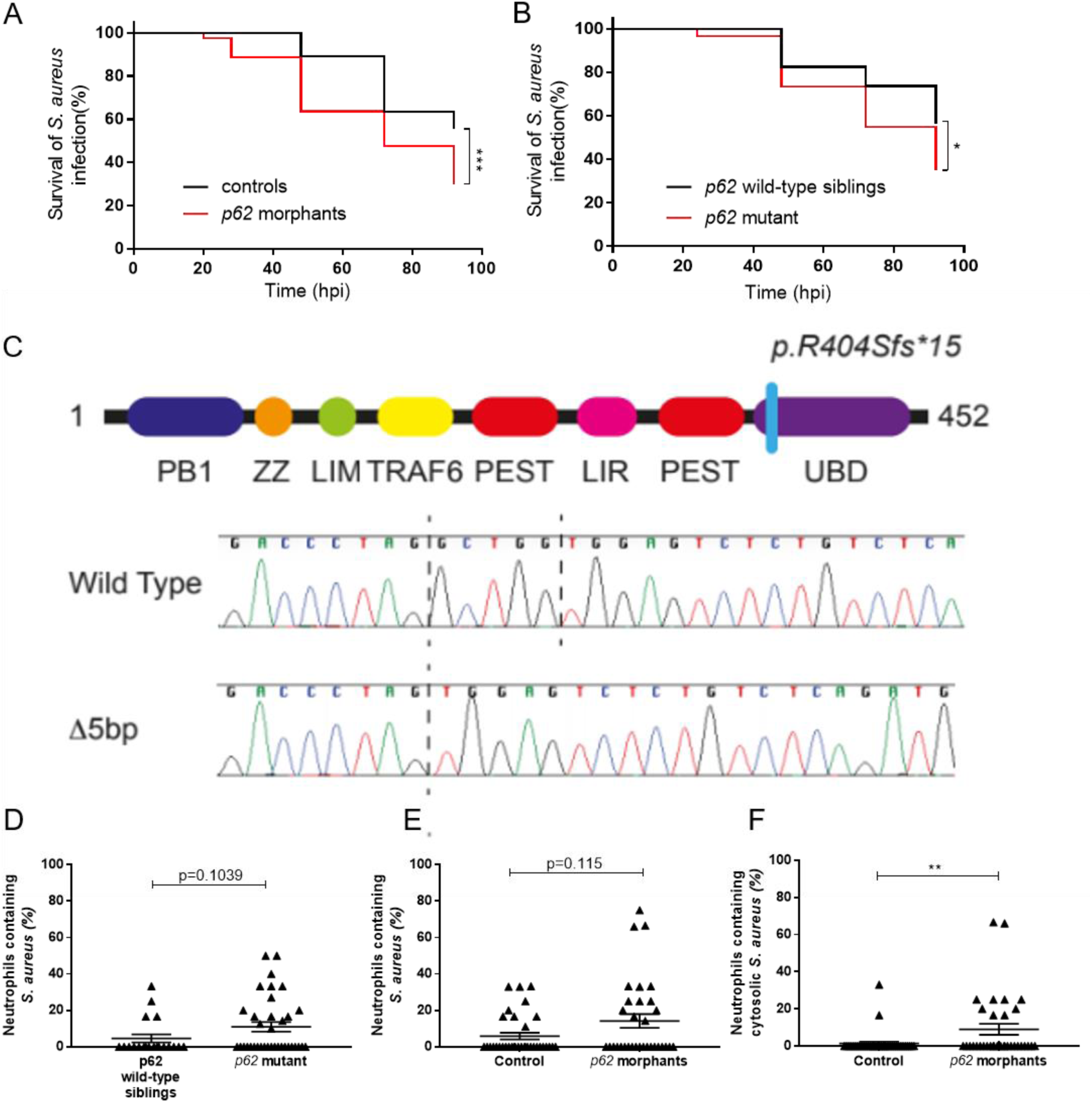
Zebrafish survival is reduced following infection with *Staphylococcus aureus* in the absence of p62. **A-B** Zebrafish survival following *S. aureus* infection, larvae were injected with 1500cfu of SH1000 at 30hpf. **A** *p62* morphants or control morphants survival (n=3, 74-80 larvae per group, p=0.004, Log-rank, Mantel-Cox test) **B** *p62* mutant or wild-type sibling survival (n=3, 57-60 larvae per group, p=0.0168, Log-rank, Mantel-Cox test) **C** Electropherograms showing the sequence of wild type and sh558 mutant p62. Dashed vertical lines show the location of the 5bp deletion. The position of the frameshift in the p62 protein is illustrated. Since this frameshift is located in the final coding exon we predict translation of a truncated p62 protein lacking the UBD domain. **D-E** Number of infected neutrophils at 26hpi following *S. aureus* infection, larvae were injected with 1500cfu of SH1000 mCherry, imaging completed in CHT (D) or GFP (E,F) at 30hpf **D** *p62* mutant or wild-type sibling (n=3, 19-36 larvae per group, p=0.0168, p=0.1039, Mann-Whitney test, error bars +/- SEM) **E** *p62* morphants or control morphants in *Tg(mpx:eGFP)*i114 larvae (n=3, 32-34 larvae per group, p=0.115, Mann-Whitney test, error bars +/- SEM) **F** *p62* morphants or control morphants *Tg(mpx:eGFP)*i114 larvae (n=3, 32-34 larvae per group, **p<0.01, Mann-Whitney test, error bars +/- SEM)

Both *p62* morpholino and *p62* mutant zebrafish (sh558) techniques do not block p62 function in neutrophils specifically, therefore we next aimed to determine whether the loss of p62 was important in neutrophils during *S. aureus* infection. Interestingly, there was no difference between the survival of our GFP-p62 reporter and wild-type controls (Supplementary Figure 4A), suggesting that endogenous *p62* expression is sufficient for restriction of the small proportion of bacteria which reside in the cytosol. First, using TSA staining of 1dpi larvae to visualise neutrophils within *p62* mutant (sh558) and control larvae, we found a non-significant (p=0.1039) increase in neutrophils containing *S. aureus* (Figure 3D). A small effect was expected due to the small proportion of cytosolic bacteria which are likely targeted by p62 during infection. It was therefore likely that showing a difference in the number of infected neutrophils would have required a very large number of infections; we were able to calculate that the observed differences would require a group size of 270. Next, using *p62* morphants and control larvae, a comparison of the number of bacteria present within neutrophils at 1dpi was completed in the *Tg(mpx:eGFP)i114 larvae*. In agreement with the p62-UBD mutant data, a non-significant (p=0.115) increase of neutrophils containing *S. aureus* was observed in *p62* morphants in comparison to wild-type controls (Figure 3E), again we were able to calculate that the observed differences would require a large group size of 219. However, examination of the bacterial location revealed a significant increase in the number of cytosolic *S. aureus* in the *p62* morphants in comparison to control fish (Figure 3F), suggesting loss of p62 is important for the control of cytosolic *S. aureus* by neutrophils. Thus, we could show that loss of *p62* leads to an increase in bacterial burden within neutrophils and that p62 is likely targeting the small proportion of bacteria which escape to the cytosol.

## Discussion

Using the unique attributes of long-term high-resolution imaging and genetic manipulation of zebrafish larvae we have shown the dynamics of Lc3 and p62 on the *S. aureus* containing vacuole, their relation to bacterial degradation, and how p62 recognises cytosolic bacteria meaning that loss of p62 activity is sufficient to increase mortality following *S. aureus* infection.

Loss of zebrafish *p62*, through morpholino-mediated knockdown, significantly increased susceptibility to infection to *S. aureus*. This is the first *in vivo* evidence that p62 is important in the outcome of intracellular handling of *S. aureus*. To confirm the *p62* knockdown data, we generated a zebrafish *p62* mutant lacking the UBD domain, which confirmed a significant increase in susceptibility to zebrafish in *S. aureus* infection. This suggests that for *S. aureus* infection control, the *p62* UBD, which is able to bind to ubiquitinated *S. aureus* (Neumann *et al.*, 2016; Singh *et al.*, 2017), is important for host control of infection. In addition to its role as an autophagy receptor, p62 can aid killing of pathogens through delivery of anti-microbial peptides (Ponpuak *et al.*, 2010); it is possible that anti-microbial peptides delivered by p62 are important in neutrophil control of *S. aureus* infection. The *p62* zebrafish mutant represents a valuable tool in the analysis of selective autophagy in infection, which may also be useful for study of other intracellular pathogens or in other diseases where autophagy is implicated in pathology, for example in neurodegenerative disorders.

Although *in vitro* studies have described co-localisation of p62 and autophagy in pathogen handling, until now, no evidence of direct p62 interactions with these pathogens has been shown in neutrophils or *in vivo*. Interaction of p62 with *S. aureus* has been demonstrated through *in vitro* studies using fibroblasts and epithelial cells (Neumann *et al.*, 2016; Singh *et al.*, 2017). *In vitro* data shows *S. aureus* can be targeted for autophagic degradation by p62 (Neumann *et al.*, 2016; Singh *et al.*, 2017), where puncta appear to be co-localised with *S. aureus*. Our new zebrafish GFP-p62 reporter shows cytosolic puncta formation, which has also been observed in other cell culture studies, both endogenous expression and using similar GFP-p62 reporter systems (Bjørkøy *et al.*, 2005; Pankiv *et al.*, 2007; Larsen *et al.*, 2010). By comparing GFP-p62 puncta marking of intracellular *S. aureus* with the location of bacteria over time, it is interesting to note that p62 marking is reduced over time for cytosolic bacteria (which appear to be a small population that persist throughout infection); this may indicate that cytosolic bacteria marked with p62 are degraded. Furthermore, at later time points in *S. aureus* infection the number of GFP-p62 puncta is reduced within infected cells, suggesting when bacteria escape the phagosome, p62 becomes important in controlling cytosolic bacteria.

We show that most *S. aureus* is contained within a vesicle soon after infection and that by 26hpi most *S. aureus* are absent from neutrophils. Of note, some images show bacteria outside neutrophils, these represent bacteria which have been phagocytosed by macrophages, which has previously been described (Prajsnar *et al.*, 2008). The large reduction of neutrophils containing bacteria from 2hpi to 26hpi, leaving a small population at 26hpi, may be representative of a niche for bacterial persistence and/or proliferation. The role of neutrophils as an intracellular niche has previously been described to be important in determining the outcome of *S. aureus* infection (Thwaites and Gant, 2011; Prajsnar *et al.*, 2012; Pollitt *et al.*, 2018). Interestingly, it appears that Lc3 marks the majority of vesicles containing bacteria. Lc3 localisation to *S. aureus* may represent Lc3 recruitment to autophagosomes, however since recruitment is observed at early infection time points, it may represent Lc3-associated phagocytosis (LAP), which is also observed in *Listeria monocytogenes* infection of macrophages (Gluschko *et al.*, 2018). Since most bacteria are degraded, it appears that Lc3 marking of vesicles could lead to bacterial degradation in the zebrafish.

Thus, we demonstrate that host p62 is beneficial for the host outcome following *S. aureus* infection and that p62 mediated control of cytosolic bacteria within neutrophils may represent one of many mechanisms employed by the host in immunity to this versatile pathogen.

## Materials and methods

### Ethics statement

Animal work was carried out according to guidelines and legislation set out in UK law in the Animals (Scientific Procedures) Act 1986, under Project License PPL 40/3574 or P1A4A7A5E). Ethical approval was granted by the University of Sheffield Local Ethical Review Panel. Animal work completed in Singapore was completed under the Institutional Animal Care and Use Committee (IACUC) guidelines, under the A*STAR Biological Resource Centre (BRC) approved IACUC Protocol #140977.

### Zebrafish husbandry

Zebrafish strains were maintained according to standard protocols (Nüsslein-Volhard and Dahm, 2002). For animals housed in the Bateson Centre aquaria at the University of Sheffield, adult fish were maintained on a 14:10-hour light/dark cycle at 28 °C in UK Home Office approved facilities. For animals housed in IMCB, Singapore, adult fish were maintained on a 14:10-hour light/dark cycle at 28°C in the IMCB zebrafish facility. LWT and AB wild-type larvae were used in addition to transgenic lines, *Tg(lyz:eGFP-p62)i330* created in this study, *Tg(lyz:RFP-GFP-Lc3)sh383* (Prajsnar *et al.*, 2019), *Tg(lyz:nfsB-mCherry)sh260* (Buchan *et al.*, 2019) (these fish encode nitroreductase gene *nsfB* within neutrophils which allows ablation of cells following metronidazole treatment, which was not used in this study) and *Tg(mpx:eGFP)i114* (Renshaw *et al.*, 2006). Generation of p62 sh558 mutant zebrafish is described below. Larvae were maintained in E3 plus methylene blue at 28°C until 5dpf.

### *S. aureus* culture

The *Staphylococcus aureus* strain SH1000 was used in this study. A single bacterial colony was placed in 10ml brain heart infusion (BHI) medium (Oxoid number 1) overnight at 37°C, 250rpm. 500μl of this overnight culture was then added to 50ml of BHI medium and incubated at 37°C, 250rpm until OD_600_ 1. The bacteria were then pelleted at 4500rpm, 4°C for 15 minutes. The bacteria were then re-suspended in PBS (Oxoid, BR0014G), using a volume to dilute to the required dose, with 1500cfu/nL being standard. Bacteria were incubated on ice for a short period, until use. Strains used: SH1000 wild-type strain (Horsburgh *et al.*, 2002), SH1000-pMV158-mCherry (Boldock *et al.*, 2018), SH1000-pMV158-GFP (Boldock *et al.*, 2018).

### Zebrafish micro-injection

For p62 morpholino micro-injections: Larvae were injected immediately after fertilisation using a *p62* morpholino (van der Vaart *et al.*, 2014). A standard control morpholino (Genetools) was used as a negative control. For injection of *S. aureus*, zebrafish larvae were injected at 1 dpf (for survival analysis, (Prajsnar *et al.*, 2008)) or 2 dpf (for microscopy analysis) and monitored until a maximum of 5dpf. Larvae were anesthetised by immersion in 0.168 mg/mL tricaine in E3 and transferred onto 3% methyl cellulose in E3 for injection. For *S. aureus* 1nl of bacteria, containing 1500cfu, was injected into the yolk sac circulation valley. Larvae were transferred to fresh E3 to recover from anaesthetic. Any zebrafish injured by the needle/micro-injection were removed from the procedure. Zebrafish were maintained at 28°C.

### Generation of *Tg(lyz:eGFP-p62)i330* transgenic line

The generation of the *Tg(lyz:eGFP-p62)i330* line was performed using the Gateway™ system in combination with Tol2 transgenesis (Tol2Kit) (Kwan *et al.*, 2007). To make the required expression clone, *pDest(lyz:eGFP-p62)*, the *p5E-lyz* entry clone (Elks *et al.*, 2011) and the *pME-eGFP-nostop* middle entry vectors were used. The destination vector *pDestTol2CG*, was chosen which included Tol2 sites for integration into the genome, in addition to a GFP heart marker. The required p62 3’ entry vector, and expression clone *pDest(lyz:eGFP-p62)* were constructed following the Multisite Gateway™ three-fragment vector construction kit. To generate Tol2 mRNA, a *pT3Tol2* plasmid was used. The DNA *pT3Tol2* plasmid was linearised through a restriction site digest. Tol2 mRNA was generated by a transcription reaction (Ambion T3 mMessage Machine) from the linear *pT3Tol2* plasmid. Tol2 mRNA and *pDest(lyz:eGFP-p62)* were co-injected into a single cell (at the single cell stage) of wild-type AB larvae. A 1nl injection contained 30pg of Tol2 mRNA and 60pg of *pDest(lyz:eGFP-p62)*.

### Microscopy of infected zebrafish

Larvae were anaesthetized 0.168 mg/mL tricaine in E3 and mounted in 0.8% low melting agarose onto glass bottom microwell dishes (MatTek P35G-1.5-14C). An UltraVIEW VoX spinning disk confocal microscope (Perkin Elmer, Cambridge, UK) was used for imaging neutrophils within larvae. 405nm, 445nm, 488nm, 514nm, 561nm and 640nm lasers were available for excitation. Most cellular level imaging was completed in the caudal hematopoietic tissue (CHT) using a 40x oil objective (UplanSApo 40x oil (NA 1.3)). In some cases a 20x objective was used for whole larvae imaging. GFP, TxRed emission filters were used and bright field images were acquired using a Hamamatsu C9100-50 EM-CCD camera. Volocity software was used. Between early and late time points zebrafish larvae were placed back into E3 and maintained at 28°C.

### pHrodo staining of *S. aureus*

Bacterial strains were prepared for injected (as above) and re-suspended into PBS pH 9. pHRODO (Thermofisher, P36600) was added at a ratio of 1:200 and incubated at 37°C for 30 minutes, shaking, in the dark. The bacteria were suspended in PBS pH 8 and washed through a series of solutions, (Tris pH8.5, PBS pH 8) and finally re-suspended into PBS pH7.4 for injection.

### Tyramide Signal Amplification (TSA) Staining

Following *S. aureus* infection, larvae were fixed in 4% paraformaldehyde overnight at 4°C. Once fixed, larvae were washed in PBS thrice. Staining of neutrophils (specifically myeloperoxidase activity) in LWT wild-type larvae was completed using TSA staining kit (Cy5-TSA Cyanine Kit, PerkinElmer, NEL705A001KT). Fish were incubated in a 1:100 ratio of Cy5-TSA:amplification diluent at 28°C for 10 minutes in the dark. Larvae were washed thrice in PBS before imaging.

### TEM of infected zebrafish

Specimens were fixed in 2.5% Glutaldehyde/0.1M Sodium Cacodylate and postfixed 2% aqueous Osmium Tetroxide, dehydrated through a graded series of ethanol, and cleared in epoxypropane (EPP) and then infiltrated in 50/50 Araldite resin: EPP mixture overnight on a rotor. This mixture was replaced with two changes over 8 hours of fresh Araldite resin mixture before being embedded and cured in a 60 degrees C oven for 48-72 hours. Ultrathin sections, approximately 85nm thick, were cut on a Leica UC6 ultramicrotome onto 200 mesh copper grids. These were stained for 10 mins with saturated aqueous Uranyl Acetate followed by Reynold’s Lead Citrate for 5mins. Sections were examined using an FEI Tecnai Transmission Electron Microscope at an accelerating voltage of 80Kv. Electron micrographs were recorded using Gatan Orius 1000 digital camera and Gatan Digital Micrograph software

### Image analysis

Image analysis was performed using ImageJ software, to quantify the number of *S. aureus* cells within neutrophils, and to quantify GFP-p62 puncta and Lc3 co-localisation to these pathogens.

### Drug treatment of zebrafish

Larvae were treated with autophagy inducers and inhibitors through immersion in E3 medium. All drugs were sourced from Sigma-Aldrich, UK. The Bay K8644 was added to the E3 to the required concentration, Bay-K 6844 1μM. Larvae were incubated at 28°C for 24 hours before microscopy. Zebrafish were not anaesthetised for immersion drug treatments.

### Generation of *p62* mutant

A zebrafish *p62* mutant was generated using CRISPR/Cas9 mutagenesis. A guide RNA targeting exon 8 of zebrafish *p62* (ACAGAGACTCCACCAGCCTA) was inserted into a published oligonucleotide scaffold (Talbot and Amacher, 2014) and injected together with recombinant Cas9 protein (New England Biolabs) into 1-2 cell stage zebrafish (AB strain). Efficiency of mutagenesis was confirmed using high resolution melt curve analysis as previously described (Sutton *et al.*, 2007) and several founders were identified. P62^sh558^ carries a 10 base pair deletion resulting in a frameshift and premature truncation of p62 in the ubiquitin-associated (UBA) domain.

### Statistical analysis

Statistical analysis was performed as described in the results and figure legends. We used Graph Pad Prism 7 (v7.04) for statistical tests and plots.

## Acknowledgments

JFG was supported by an award from the Singapore A*STAR Research Attachment Programme (ARAP) in partnership with the University of Sheffield, and a Medical Research Council Grant (MR/R001111/1 with SAR and SJF). TKP was supported by an individual Marie Curie fellowship (PIEF-GA-2013-625975) and by AMR cross-council funding from the MRC to the SHIELD consortium “Optimising Innate Host Defence to Combat Antimicrobial Resistance” MRNO2995X/1. RDT and AJG were supported by the University of Sheffield. JJS was a Marie Curie fellow in the Initial Training Network FishForPharma (PITN-GA-2011-289209). Work in the PWI lab was funded by the A*STAR Institute of Molecular and Cell Biology (IMCB) and the Lee Kong Chian School of Medicine. SAJ was supported by Medical Research Council and Department for International Development Career Development Award Fellowship MR/J009156/1 (http://www.mrc.ac.uk/). SAJ was additionally supported by a Krebs Institute Fellowship (http://krebsinstitute.group.shef.ac.uk/), and Medical Research Council Centre grant (G0700091). SAR was supported by a Medical Research Council Programme Grant (MR/M004864/1) (http://www.mrc.ac.uk/). Imaging was completed at the Wolfson Light Microscopy Facility. We thank aquarium staff at the Bateson Centre (Sheffield) and the IMCB (Singapore) for zebrafish husbandry.

## Author contributions

JFG, TKP, SAJ, AJG, PWI and SAR conceived this study and designed and interpreted experiments. JFG prepared manuscript with important intellectual input from TKP, SAJ, AJG, SJF and SAR. JFG and JJS conducted bacterial fate and location analysis. JFG performed GFP-p62 reporter line generation and characterisation, zebrafish infection, morpholino experiments, intracellular imaging and subsequent analysis. AKT performed zebrafish infection. CJH performed TEM. SAJ performed imaging of bacteria degradation. TKP performed p62 morpholino experiments. AJG and RDT generated the p62 stable mutant line.

---

**Supplementary Figure 1:**
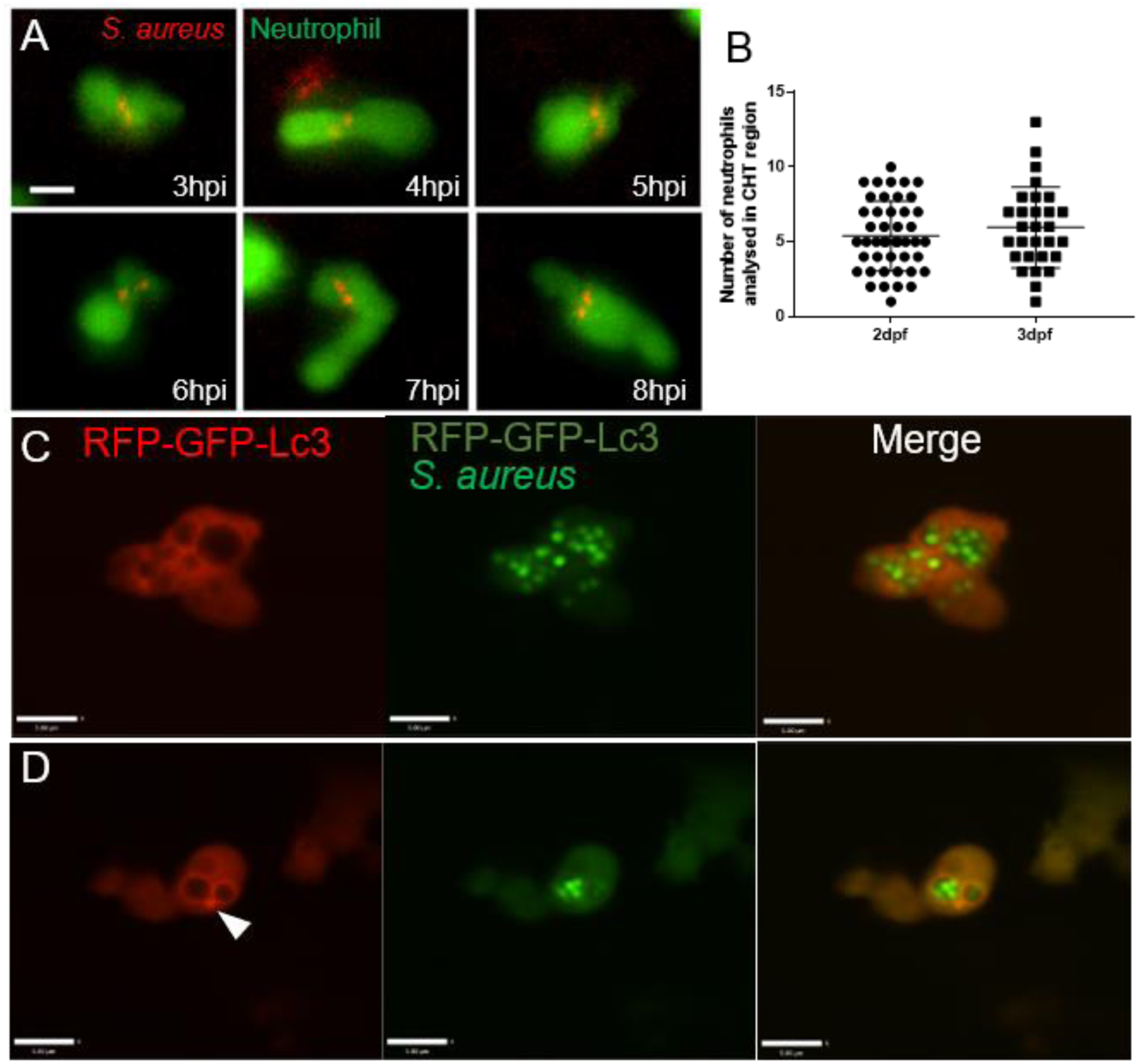
Neutrophil interactions with *S. aureus* in the zebrafish model. **A** *Tg(mpx:eGFP)*i114 larvae were injected at 1dpf with 1500cfu SH1000-mCherry *S. aureus*, and imaged at 3 hours post infection. Images were captured every 5 mins for 12 hours at multiple z planes to follow infected neutrophils over time. Scale bar is 5μm. **B** Neutrophil counts in CHT region in 2dpf fish and 3dpf used for analysis (n=6, 29-45 larvae per time point, no significant difference, unpaired T-test) **D-E** *Tg(lyz:RFP-GFP-Lc3)sh383* larvae were injected at 2dpf with GFP *S. aureus*, and imaged at 2hpi, and ∼26hpi. **C** *S. aureus* without Lc3 marking the entire vesicle, scale 5.8um. Also demonstrating a clear vesicle structure **D** *S. aureus* with Lc3 puncta marking the edge of a vesicle, (indicated with an arrowhead), scale 9μm

**Supplementary Figure 2:**
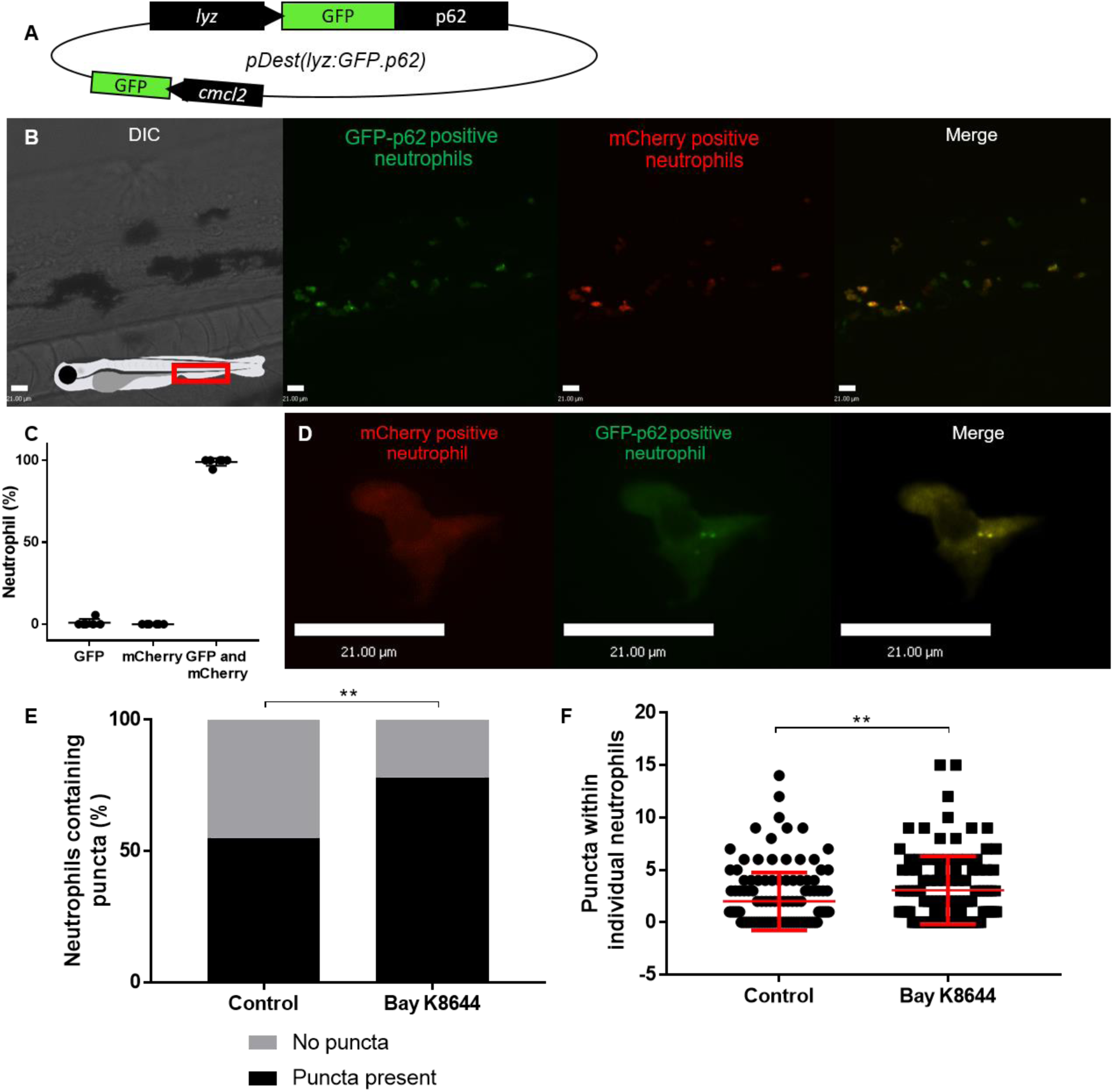
The GFP-p62 reporter line expresses a GFP-p62 fusion protein within neutrophils. **A** Illustration of the plasmid created to make the GFP-p62 zebrafish reporter line, pDest(lyz:eGFP*-p62*). **B** Representative image of neutrophils within larva resulting from crossing *Tg(lyz:nfsB-mCherry)sh260* (which has mCherry fluorescent neutrophils), with *Tg(lyz:eGFP-p62)i330* which shows GFP expression within neutrophils. Image taken of the caudal haematopoietic tissue (CHT) shown in red box of diagram of larvae. **C** The percentage of neutrophils which express fluorescence of GFP alone, mCherry alone, or both GFP and mCherry using six representative larvae from a cross of *Tg(lyz:nfsB-mCherry)sh260* with *Tg(lyz:eGFP-p62)i330* zebrafish lines (CHT imaged, one-way ANOVA, Tukey’s multiple comparisons ****=p<0.001). **D** High magnification imaging of an individual neutrophil from the CHT showing GFP puncta, but not mCherry puncta, from a *Tg(lyz:nfsB-mCherry*)*sh260* cross with *Tg(lyz:eGFP-p62)i330*. **E-F** Quantification of images taken of *Tg(lyz:eGFP-p62)i330* larvae in the CHT, treated with autophagy blocking drug Bay K8644. **E** Percentage of neutrophils with puncta present (Chi-square test, **p<0.01, n=3, 15-20 larvae per group) **F** Number of puncta observed within individual neutrophils (Error bars +/- SD, Unpaired T-test **p<0.01, n=3, 15-20 larvae per group)

**Supplementary Figure 3:**
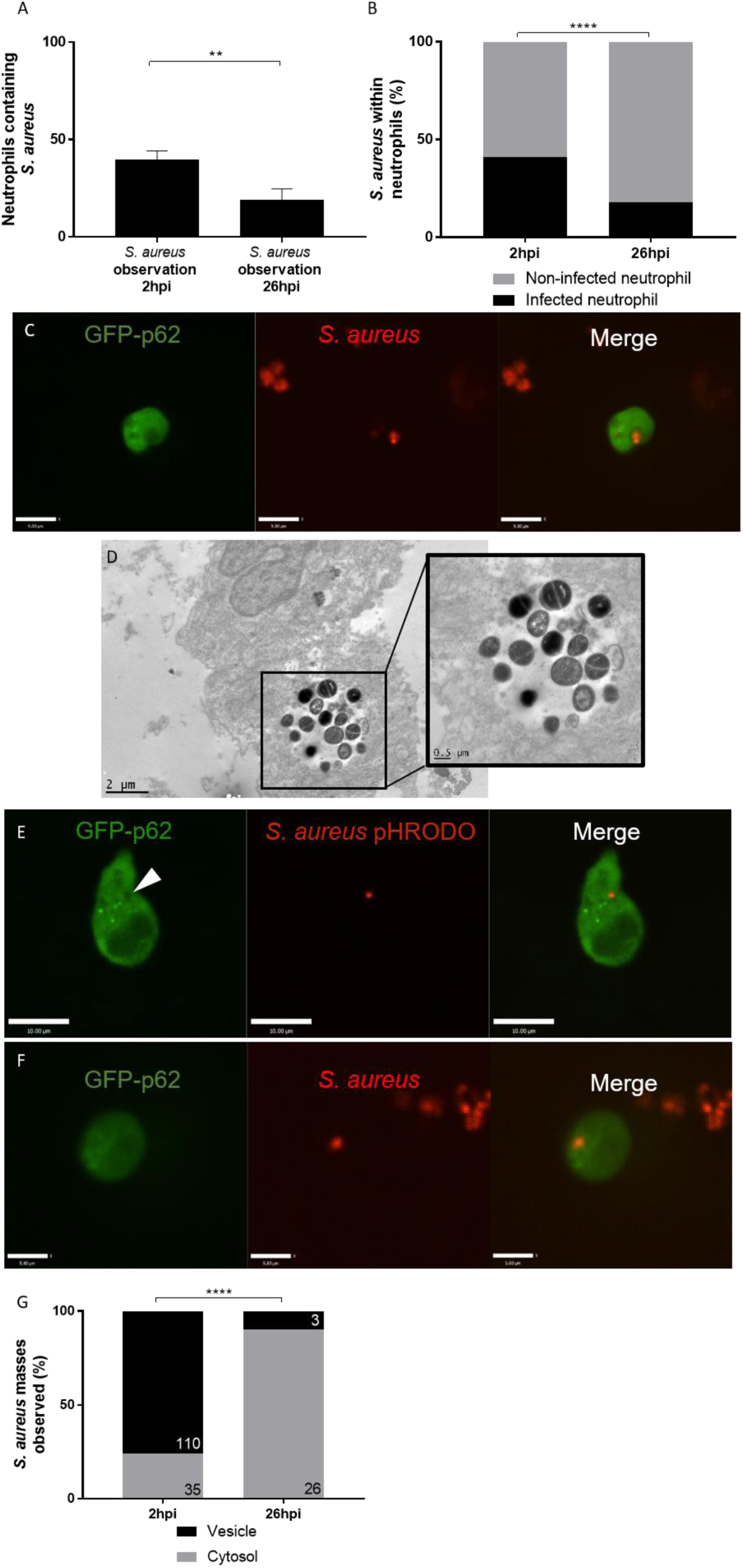
The intracellular location of *Staphylococcus aureus* within neutrophils changes throughout infection. **A-C and F-G** *Tg(lyz:eGFP-p62)i330* larvae were injected with mCherry *S. aureus* at 2dpf, and imaged at 2hpi, and ∼26hpi in the CHT. **A** The number of *S. aureus* events observed within neutrophils at 2hpi and 26hpi (n=3, 27 larvae at 2hpi, and 18 larvae at 26hpi unpaired t-test, **p<0.01, +/- SEM) **B** Proportion of neutrophils infected or non-infected with *S. aureus* at 2hpi and 26hpi (****p<0.001, Fisher’s exact test, n=3, 14 larvae at 2hpi, and 12 larvae at 26hpi) **C** Representative image of *S. aureus* observed within a likely “vesicle” **D** LWT zebrafish larvae injected with 1500cfu of SH1000 *S. aureus* and fixed at ∼18hpi. Transmission electron micrograph of an infected neutrophil. Highlighted box shows zoom in of the vesicle contain *S. aureus*. **E** *Tg(mpx:eGFP)*i114 larvae injected at 1dpf with 1500cfu SH1000 *S. aureus* stained with pHrodo and imaged at 2hpi imaged in the CHT **F** representative image of *S. aureus* observed within the cytosol **G** Proportion *S. aureus* events observed within vesicles or cytosol at 2hpi and 26hpi (****p<0.001, Fisher’s exact test, n=3, 14 larvae at 2hpi, and 12 larvae at 26hpi)

**Supplementary Figure 4:**
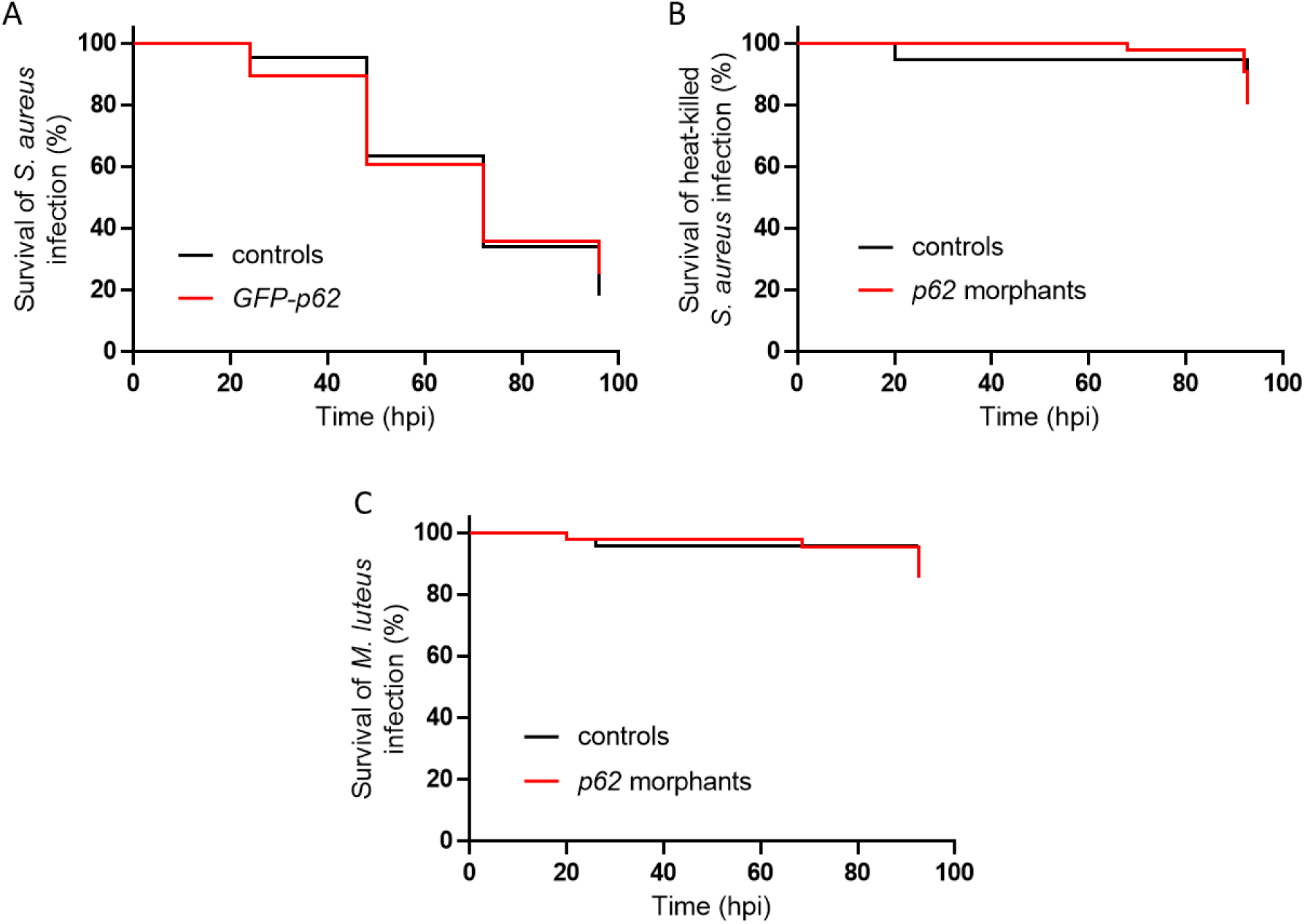
The role of p62 in zebrafish survival following infection. **A** Zebrafish survival following *S. aureus* infection,GFP-p62 larvae or controls were injected with 1500cfu of SH1000 at 30hpf (n=3, 39-44 larvae per group, Log-rank, Mantel-Cox test) **B** Zebrafish survival following injection of heat-killed *S. aureus* (equivalent of 1500cfu SH1000) at 30hpf, *p62* morphants or control morphants (n=3, 38-44 larvae per group, Log-rank, Mantel-Cox test) **C** Zebrafish survival following injection of *M. luteus* 2000cfu at 30hpf, *p62* morphants or control morphants (n=3, 44-45 larvae per group, Log-rank, Mantel-Cox test)

## Supplementary video legends

**Supplementary Video 1: GFP-p62 puncta co-localise with a vesicle containing *S. aureus***

*Tg(lyz:eGFP-p62)i330* larvae were injected with 1500cfu mCherry *S. aureus* at 2dpf and imaged at 3hpi in the CHT. Images were collected as fast as possible (∼every 0.8 seconds) for up to 3 minutes. White arrow indicates the location of the vesicle marked with GFP-p62 puncta.

**Supplementary Video 2: GFP-p62 puncta co-localise with *S. aureus* located in the cytosol**

*Tg(lyz:eGFP-p62)i330* larvae were injected with 1500cfu mCherry *S. aureus* at 2dpf and imaged at 3hpi in the CHT. Images were collected as fast as possible (∼every 0.8 seconds) for up to 3 minutes. White arrow indicates the location of the S. aureus in the cytosol marked with GFP-p62 puncta.

